# SHIV.CH505-infected infant and adult rhesus macaques exhibit similar HIV Env-specific antibody kinetics, despite distinct T-follicular helper (Tfh) and germinal center B cell landscapes

**DOI:** 10.1101/538876

**Authors:** Ashley N. Nelson, Ria Goswami, Maria Dennis, Joshua Tu, Riley J. Mangan, Pooja T. Saha, Derek W. Cain, Xiaoying Shen, George M. Shaw, Katharine Bar, Michael Hudgens, Justin Pollara, Kristina De Paris, Koen K.A. Van Rompay, Sallie R. Permar

**Affiliations:** Human Vaccine Institute, Duke University Medical Center, Durham, NC, USA; Gillings School of Public Health and Center for AIDS Research, University of North Carolina at Chapel Hill, Chapel Hill, North Carolina, USA; Department of Medicine, University of Pennsylvania, Philadelphia, PA, USA; Department of Microbiology and Immunology and Center for AIDS Research, School of Medicine, University of North Carolina at Chapel Hill, Chapel Hill, North Carolina, USA; California National Primate Research Center, University of California, Davis, CA, USA

**Author notes:** Address correspondence to Sallie R. Permar,. A.N.N. and R.G. contributed equally to this work.

## Abstract

Pediatric HIV infection remains a large global health concern despite the widespread use of antiretroviral therapy (ART). Thus, global elimination of pediatric HIV infections will require the development of novel immune-based approaches, and understanding infant immunity to HIV is critical to guide the rational design of these intervention strategies. Despite their immunological immaturity, HIV-infected children develop broadly neutralizing antibodies (bnAbs) more frequently and earlier than adults do. Furthermore, T-follicular helper (Tfh) cells have been associated with bnAb development in HIV-infected children and adults. To further our understanding of age-related differences in the development of HIV-specific immunity, we evaluated the generation of virus-specific humoral immune responses in infant (n=6) and adult (n=12) rhesus macaques (RMs) infected with a transmitted/founder (T/F) simian-human immunodeficiency virus (SHIV.C.CH505). The plasma HIV envelope-specific IgG antibody kinetics were similar in SHIV-infected infant and adult RMs, with no significant differences in the magnitude or breadth of these responses. Interestingly, autologous tier 2 virus neutralization responses also developed with similar frequency and kinetics in infant and adult RMs, despite infants exhibiting significantly higher Tfh and germinal center B cell frequencies compared to adults. Our results indicate that the humoral immune response to SHIV infection develops with similar kinetics among infant and adult RMs, suggesting that the early life immune system is equipped to respond to HIV-1 and promote the production of neutralizing HIV antibodies.

**Importance:** There is a lack of understanding on how the maturation of the infant immune system influences immunity to HIV infection, or how these responses differ from those of adults. Improving our knowledge of infant HIV immunity will help guide antiviral intervention strategies that take advantage of the unique infant immune environment to successfully elicit protective immune responses. We utilized a rhesus macaque model of SHIV infection as a tool to distinguish the differences in HIV humoral immunity in infants versus adults. Here, we demonstrate that the kinetics and quality of the infant humoral immune response to HIV are highly comparable to that of adults during the early phase of infection, despite distinct differences in their Tfh responses, indicating that slightly different mechanisms may drive infant and adult humoral immunity.

## Introduction

Despite the widespread availability of anti-retroviral (ARV) therapy for HIV-infected pregnant women, approximately 180,000 infants were newly infected in 2017 due to issues of treatment access and adherence, and acute maternal HIV infection (1). While the developing immune system and lack of immunological memory during early infancy can render neonates more susceptible to infections, it may also provide an opportunity for unique interventions. Furthermore, HIV immunization in infancy could be an opportunity to both interrupt postnatal transmission to breastfeeding infants, the current most common mode of infant HIV infection (2), as well as elicit life-long HIV immunity prior to the renewed HIV acquisition risk upon sexual debut. A better understanding of how the infant’s developing immune system influences disease outcome and pathogenesis during HIV infection and how it compares to adults is imperative to inform the design and evaluation of pediatric intervention therapies and vaccination strategies.

The disease course in HIV-infected infants is dramatically different from that of HIV-infected adults (3). Without treatment, vertically HIV-infected infants tend to have higher plasma viral RNA loads, experience rapid declines in peripheral CD4+ T cell counts, and progress to AIDS more rapidly than adults (4, 5). However, transmission studies have indicated that infants infected during breast-feeding tend to have a better clinical outcome. Specifically, postnatal HIV-infected infants have a lower risk of mortality within the first 18 months of infection (6), increased median survival times from infection (7), and better long-term survival rates (8, 9) than infants who acquire HIV infection perinatally. The ontogeny of HIV Env-specific antibodies is also quite different between infants and adults, and understanding these differences could inform infant HIV Env vaccine development and evaluation. In HIV-infected adults, Env-specific antibodies are detectable by approximately 14 days after infection (10). Yet the early kinetics of HIV-exposed, infected infants’ natural IgG responses are masked by placentally-acquired maternal antibody. More importantly, increases in HIV-specific IgG responses from 6 months through the first year of life have been implicated in a better clinical outcome (11, 12). Interestingly, recent studies have indicated that HIV-infected infants frequently develop broadly neutralizing antibodies (bnAbs) during early life (range: 11.4-28.2 months), and a bnAb isolated from an infected infant exhibited lower levels of somatic hypermutation than adult-isolated bnAbs with similar potency and breadth (13-15). Despite the early development of bnAbs in infants, the presence of antibodies capable of mediating antibody-dependent cellular cytotoxicity (ADCC) have been reported to be delayed in infants infected during the first 6 weeks of life, yet are associated with a better outcome of disease (16, 17). Thus, it is likely that the immune mechanisms associated with disease progression in pediatric HIV differ from those of adults, and further studies in animal models would help determine key features of the infant immune landscape that have the greatest influence on disease pathogenesis.

Experimental infection of non-human primates (NHP) with chimeric simian-human immunodeficiency virus (SHIV) remains an invaluable model for studying HIV pathogenesis, and evaluating therapeutic and prevention strategies. In addition, the rhesus macaque-SHIV model can be used to define differences in the ontogeny of immune responses directed against HIV between infants and adults. We have utilized this model to study adult and infant humoral immune responses to acute infection with SHIV.C.CH505, a next generation SHIV expressing the Env glycoprotein of the HIV-1 transmitted/founder CH505 (subtype C) virus, isolated from a HIV-infected adult that developed plasma bnAb activity (18-20). A mutation within the CD4 binding site of this SHIV Env facilitates enhanced interaction with the rhesus CD4 molecule (18). More importantly, SHIV.C.CH505 can infect and replicate efficiently in rhesus macaques, and exhibit plasma viral load kinetics in macaques similar to those seen in acute HIV-1 infections in humans (18). Using this highly relevant infection model, we conducted a longitudinal analysis of the virus-specific humoral immune response in SHIV.C.CH505-infected infant and adult monkeys, and defined the virologic and immunologic features that are associated with the development of plasma neutralization activity in each age group. This work will allow us to improve our understanding of age-related differences in HIV Env-specific B-cell immunity elicited during acute infection, which can guide the development of infant vaccine strategies that can optimally prime B cells for virus neutralizing responses.

## Results

### Infant and adult RMs exhibit similar plasma viral load kinetics following SHIV.C.CH505 infection

Twelve female, adult rhesus macaques (age range: 4-10 y.o; Table 1) were infected intravenously with SHIV.C.CH505 at a dose of 3.4×10^5^ TCID_50_. All twelve adult monkeys became infected after the first inoculation. To mimic breast milk transmission, six infant rhesus macaques were infected orally starting at 4 weeks of age (Table 1) with a repeated exposure regimen that has been successfully developed with SIV_mac251_ to simulate exposure to HIV by breastfeeding (21, 22). Once infected, viremia peaked at 2 weeks in both age groups and the peak viremia was of similar magnitude between the groups (infant range: 6.7×10^5^−3.2 x10^7^; adult range: 3 x10^5^−1.3 x10^7^vRNA copies/mL of plasma) (Fig. 1A). At 12 wpi, 11 of the 12 adult monkeys, and 4 of the 6 infant monkeys maintained plasma viral loads above the limit of detection of the assay (15 vRNA copies/mL of plasma) (Fig. 1A). Both infant and adult monkeys exhibited decreased frequencies of CD4+ T cells at 3 wpi (Fig. 1B), however these frequencies were maintained through 12 wpi with no observed changes in CD4+ T cell counts (Fig. 1C), thus the SHIV.C.CH505 virus exhibited a more attenuated phenotype in infant monkeys in contrast to observations in the SIV_mac251_ model (23-25). Since the same viremic pattern was observed in both age groups independent of the route of infection, this model provided an opportunity to define differences in the infant and adult immune response to SHIV infection.

**Table 1.**
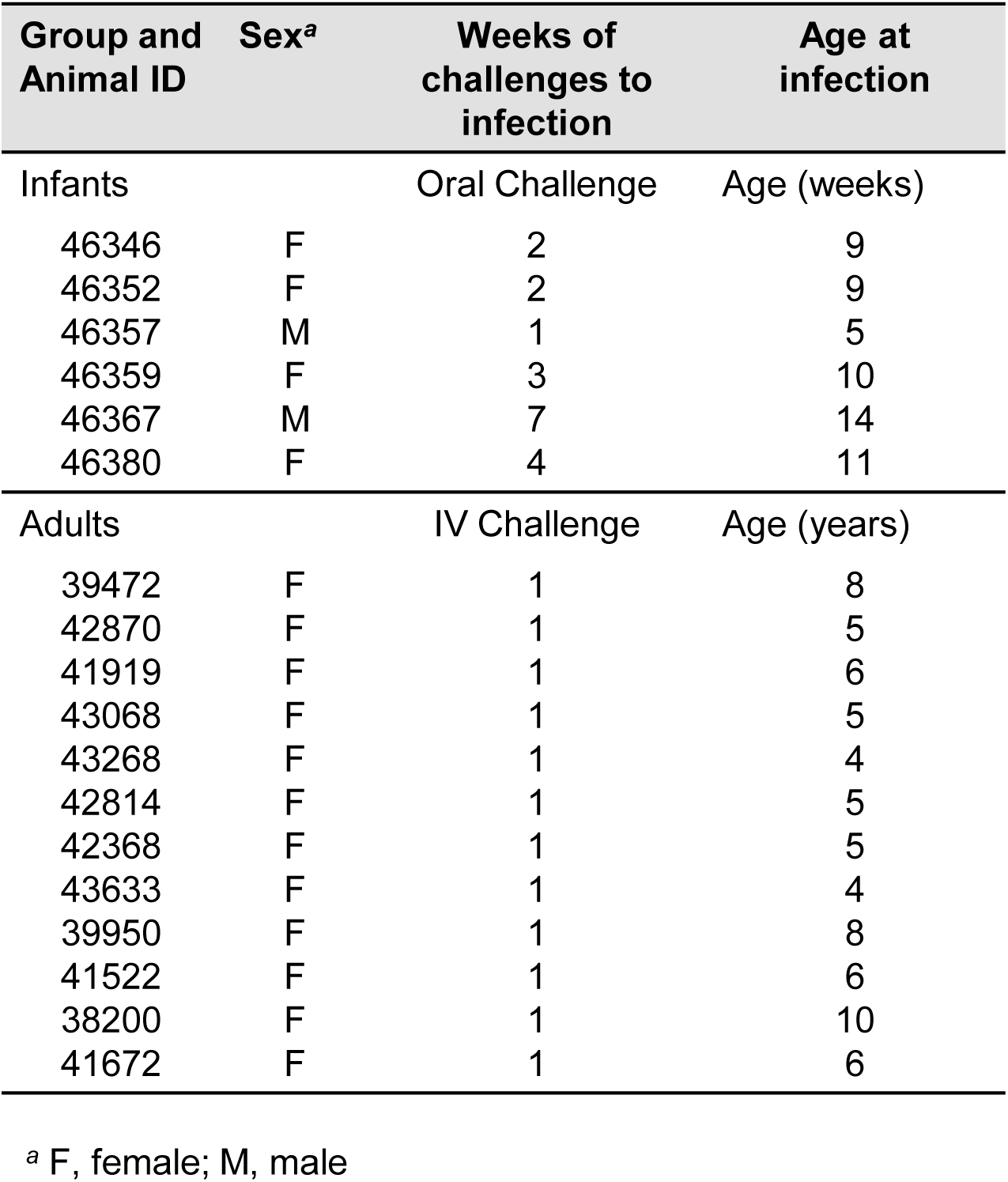
Infant and adult SHIV.C.CH505-infected monkey cohort information, number of challenges to infection, and age at infection

**Figure 1.**
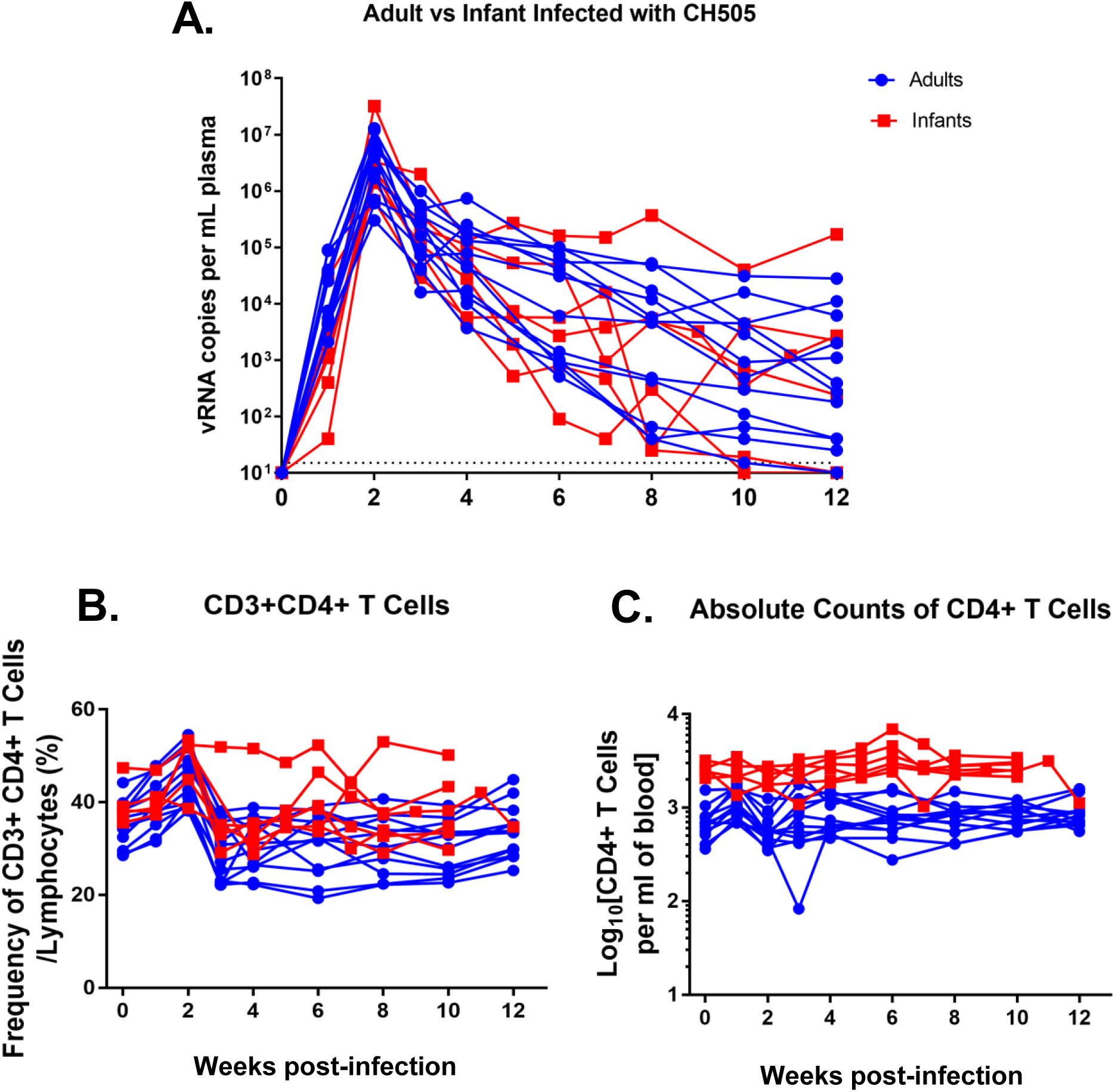
Plasma viral load and CD4+ T cell frequencies adult and infant monkeys following SHIV.C.CH505 infection. (A) Plasma viral loads were monitored weekly or bi-weekly through 12 wpi. Automated complete blood counts were collected weekly and (B) Proportion and (C) absolute counts per ml of blood of CD4+ T cells were determined. Blue lines represent adult monkeys, while red lines represent infant monkeys. Each line represents one animal.

### The kinetics, magnitude, and specificity of HIV Env-specific plasma IgG responses are similar in infant and adult RMs during acute SHIV.C.CH505 infection

We measured HIV Env-specific antibody responses to evaluate whether the ontogeny of these responses would differ between the infant and adult monkeys. In both groups, CH505 gp120-specific responses were detectable by 3 wpi with no statistically significant difference in magnitude (median gp120 IgG in infants and adults, 1,297 ng/ml and 1785 ng/ml, respectively, p=0.301) (Fig. 2A). These responses continued to increase through 12 wpi with no significant difference in magnitude (median gp120 IgG in infants and adults, 1.3x 10^5^ ng/ml and 2.3×10^5^ ng/ml, respectively, p=0.494) or kinetics between infants and adults (Fig. 2A). Overall, the kinetics and magnitude of the gp41-specific plasma IgG response was also not significantly different between both age groups (Fig. 2B). However, at 12 wpi the infant monkeys exhibited a trend towards lower gp41-specific IgG responses compared to that of adults, although this difference was not significant after correction for multiple comparisons (median gp41 IgG in infants and adults at 12 wpi, 3,712 ng/ml and 23,305 ng/ml, respectively, p=.041; FDR p=.458) (Fig. 2B).

**Figure 2.**
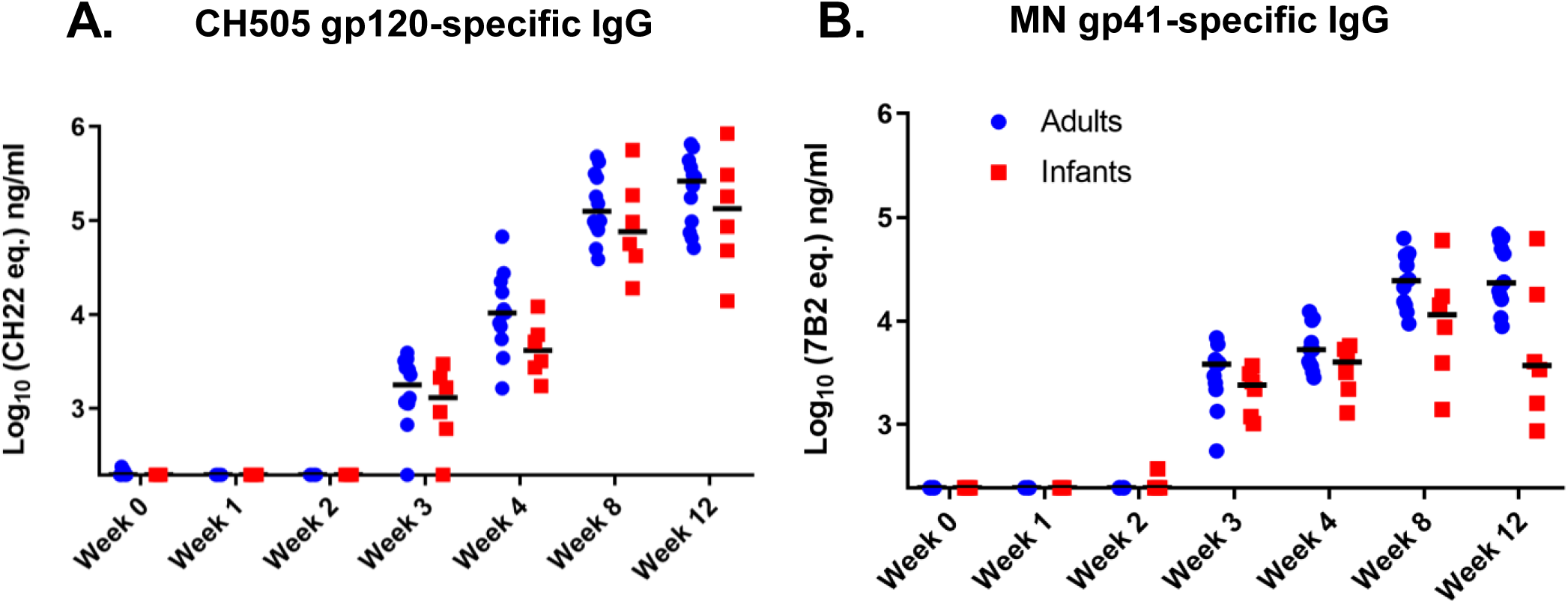
The kinetics and magnitude of HIV Env-specific IgG responses are similar in infant and adult monkeys during acute SHIV.C.CH505 infection. HIV CH505 gp120-(A) and MN gp41-specific (B) IgG responses in the plasma of adult (blue circles) and infant (red squares) monkeys through 12 wpi are shown. Statistical analysis was performed using Wilcoxon rank sum tests with exact p-values to compare IgG responses between SHIV-infected infant and adult monkeys, followed by adjustments for multiple comparisons. *unadjusted p<0.05. All p-values are >0.05 once adjusted for multiple comparisons (See Table S3 for both unadjusted p and FDR_p for all comparisons) Medians are indicated as black horizontal lines on the dot plots.

To determine the specificity of the plasma Env-specific IgG responses between age groups, we used a binding antibody multiplex assay (BAMA) to assess binding to various HIV Env linear and conformational epitopes at weeks 4 (Fig. 3A) and 12 (Fig. 3B) post-infection. For both groups, anti-V3 and –C5 binding responses were dominant and increased from week 4 to week 12 (Fig. 3A and B). Although infant monkeys exhibited a slightly higher antibody specificity for the CD4 binding site at 3 wpi, these differences were not statistically significant (p=0.438; Fig. 3C). While anti-V2 responses were rarely detected by BAMA (Fig. 3A and B), linear peptide microarray analysis demonstrated that these responses were primarily CH505-specific (Fig. S1). We also used BAMA to assess cross-clade gp120 and gp140 breadth at 4 and 12 wpi (Fig. 4A and B). At 12 wpi, antibodies from infants and adults recognized all nine gp120 and gp140 antigens tested demonstrating breadth acquisition (Fig 4B). Overall, the median gp120- and gp140-specific IgG responses across all clades trended higher at week 12 in adult monkeys, consistent with a higher gp41 plasma IgG binding response (Fig. 4B).

**Figure 3.**
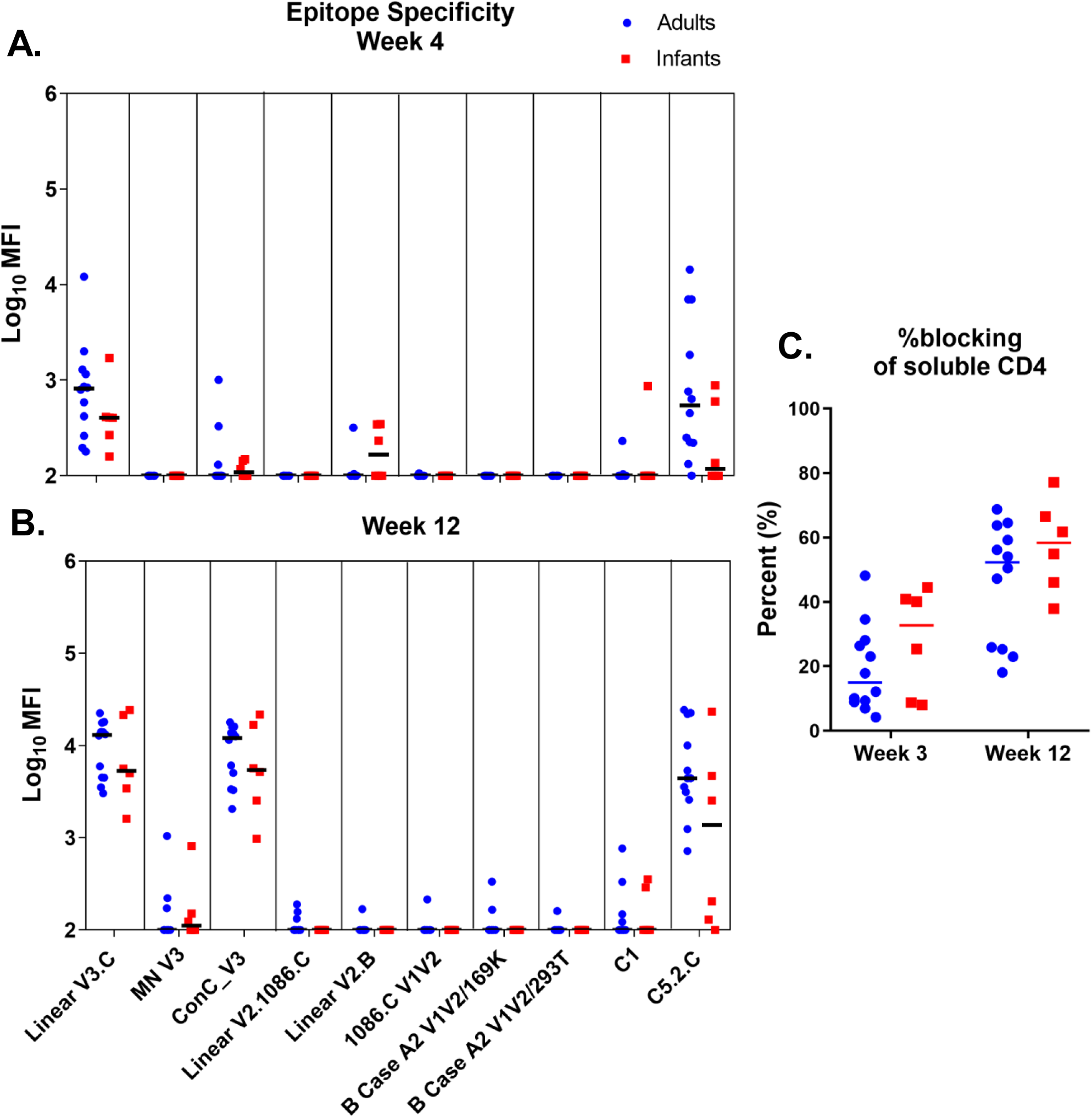
Similar specificity of Env-specific IgG responses during acute SHIV.C.CH505 infection in infants and adults. Plasma IgG specificity against a panel of HIV Env linear and conformational epitopes at week 4 (A) and week 12 (B) post-infection. (C) Plasma blocking of soluble CD4-gp120 interactions at week 3 and 12 post-infection. Adult monkeys are represented by blue circles, and infant monkeys are represented by red squares. Medians are indicated as horizontal lines on the dot plots.

**Figure 4.**
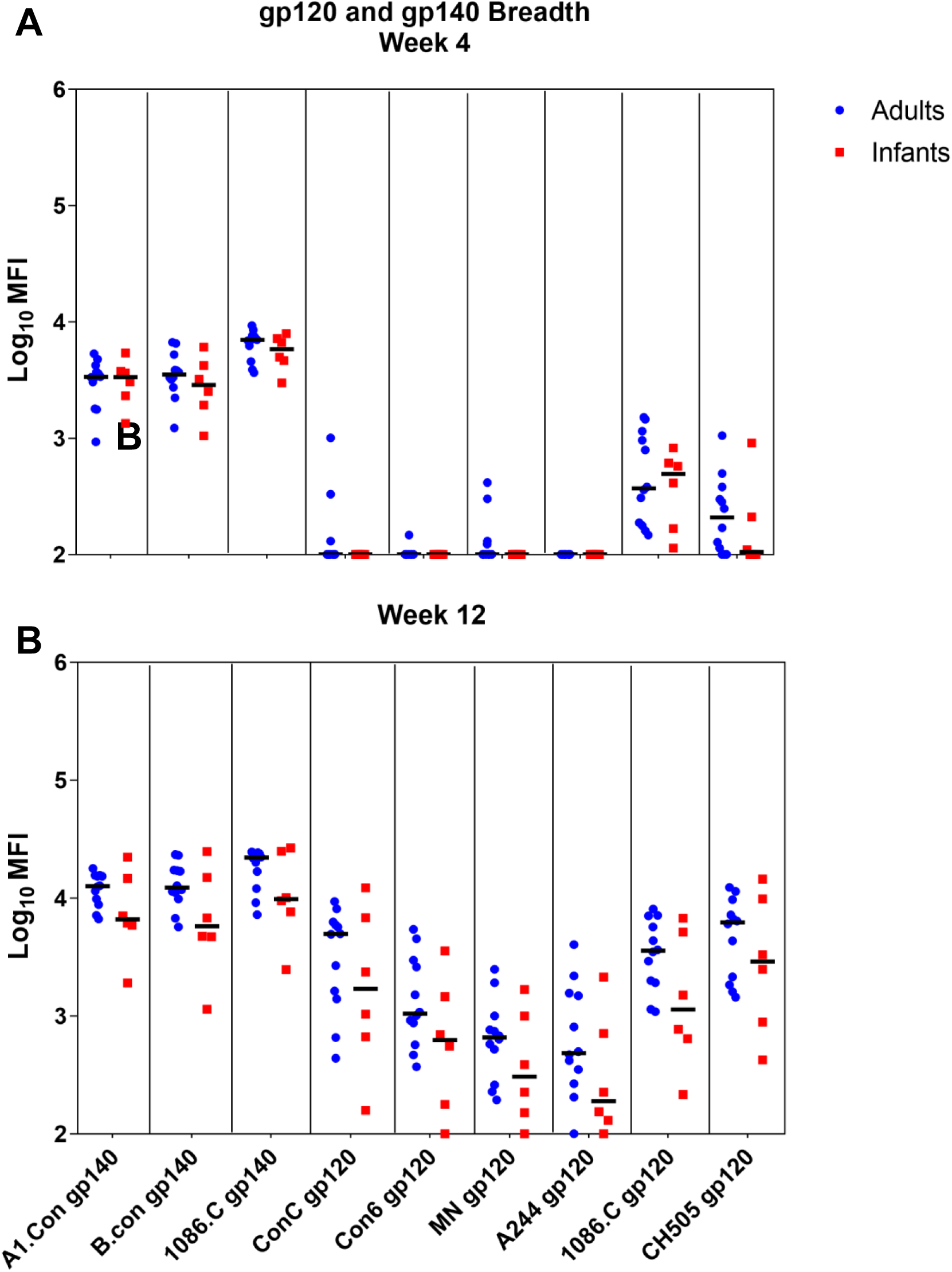
Adult and infant monkeys developed a similar breadth in gp120 and gp140 IgG responses during acute SHIV.C.CH505 infection. Cross-clade HIV gp120 and gp140 IgG breadth at week 4 (A) and week 12 (B) post-infection. Adult monkeys are represented by blue circles, and infant monkeys are represented by red squares. Medians are indicated as horizontal lines on the dot plots.

### SHIV.C.CH505-infected infant monkeys have higher proportions of T-follicular helper (Tfh) cells in the lymph node compared to that of adults

Frequencies of Tfh cells have been reported to be increased in HIV-infected children compared to adults (26). In order to investigate Tfh cell responses during the early phase of SHIV infection, we evaluated the proportions of CH505-specific (Fig. 5A) and total CD4+ CXCR5^hi^PD1^hi^(Fig. 5B) Tfh cells in the lymph node of infant and adult monkeys at 12 wpi. While the proportion of CH505-specific Tfh cells was not significantly different between the two groups (p=0.592), the infants had significantly higher proportions of total CD4+ CXCR5^hi^PD1^hi^Tfh cells (p=0.024; FDR p=.036) (Fig. 5A and B). Additionally, we compared the frequencies of the Tfh subsets based on the surface expression of CXCR3 and CCR6 as follows: Tfh1 (CXCR3+ CCR6-), Tfh2 (CXCR3-CCR6-), and Tfh17 (CXCR3-CCR6+) (Fig. 5C-E). In all, adult monkeys exhibited a significantly higher proportion of Tfh1 cells in the lymph node (p= 0.013; FDR p=.021) (Fig. 5C), while infants had a significantly higher proportion of Tfh17 cells (p= 0.001; FDR p=.003) (Fig. 5E) at 12 wpi. No significant difference was observed in Tfh2 proportions between the two age groups (p=0.066) (Fig. 5D).

**Figure 5:**
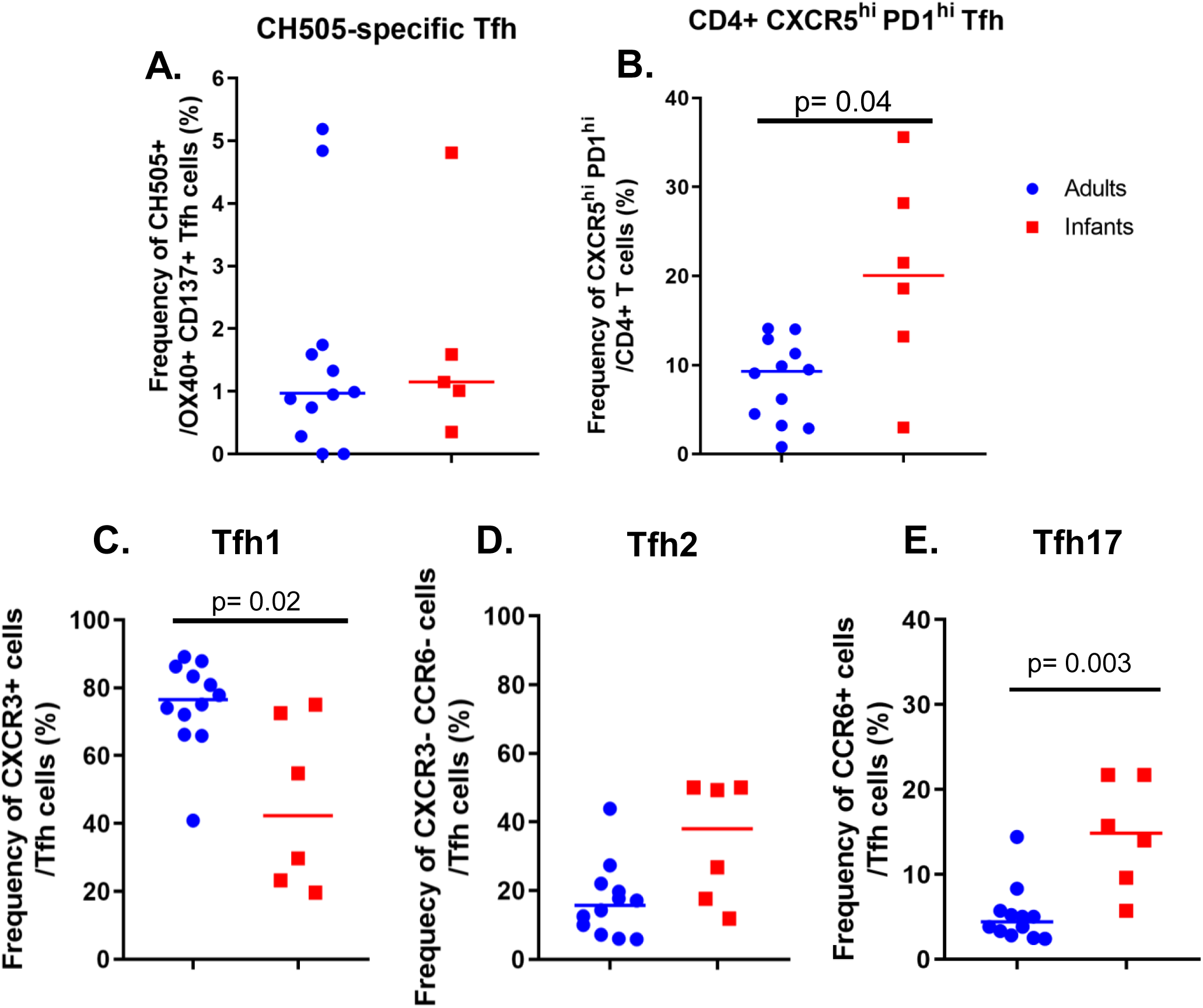
Frequency of Follicular T helper cells (Tfh) in the lymph node of SHIV.C.CH505-infected infant and adult monkeys at 12 wpi. Proportions of (A) CH505-specific and (B) CXCR5^hi^PD1^hi^Tfh cells. Proportions of Tfh subsets (C) CXCR3+ Tfh1, (D) CXCR3-CCR6-Tfh2, and (E) CCR6+ Tfh17 cells. Each data point represents one animal, and medians are indicated as horizontal lines. FDR adjusted p-values are reported in the graphs, FDR_p <0.05 was considered significant. See Tables S3 and S4 for both unadjusted p and FDR_p for all comparisons.

### Proportions of CH505-specific memory B cells are similar between infant and adult monkeys

We evaluated systemic memory B cells in infant and adult monkeys at 0, 6, and 12 wpi (Fig. 6A and B). Additionally, we compared the frequency of total B cells and germinal center (GC) B cells in the lymph node at 12 wpi between both age groups (Fig. 6C-E). Due to lack of sample availability, memory B cell populations at week 0 in the systemic compartment were only evaluated in 3 monkeys from each age group. Changes from baseline in total memory B cells (CD14-CD16-CD20+ IgD-CD27+) and CH505 gp120-specific memory B cells, were not significantly different between age groups at 6 and 12 wpi (p=1; 6 and 12 wpi for both parameters) (Fig. 6A and B). Similarly, the frequencies of total B cells (CD3-CD20+) in lymph nodes were not significantly different (p = 0.143) (Fig. 6D) at 12 wpi. However, infants exhibited a significantly higher frequency of GC B cells (CD20+ Bcl6+ Ki67+) (Fig. 6C) compared to adults (p= 0.03; FDR p=0.05) (Fig. 6E), consistent with high proportions of Tfh cells in the lymph node at 12 wpi (Fig. 5B).

**Figure 6:**
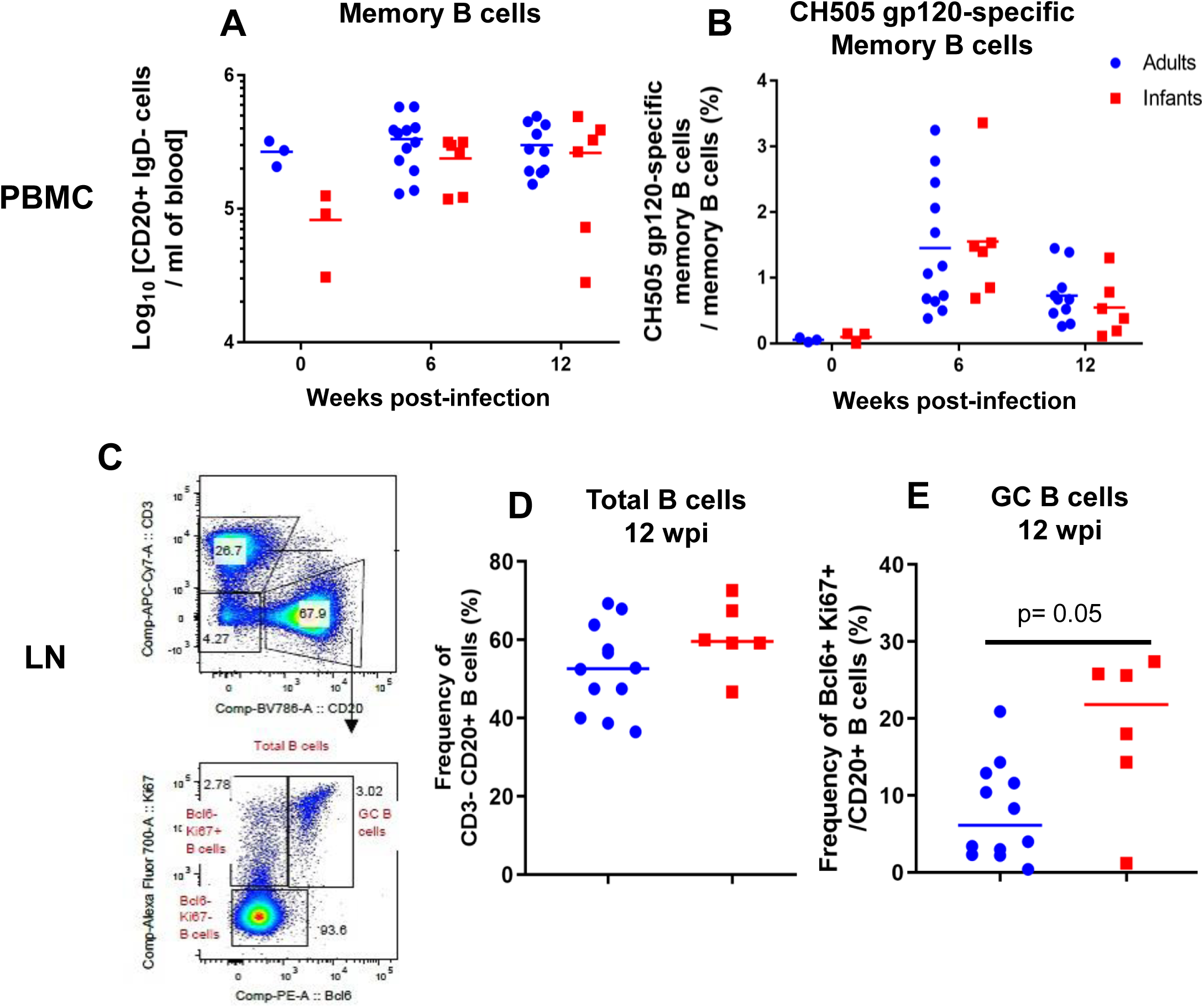
Similar proportion of systemic and lymph node B cell subsets in SHIV.C.CH505-infected infant and adult RMs. (A) Absolute counts of memory B cells (CD14-CD16-CD20+ IgD-CD27 all)/ ml of blood and (B) frequency of CH505 gp120-specific memory B cells of total memory B cells. GC B cells were identified as CD20+ BCL6+ Ki67+ cells (C), and the frequency of (D) CD3-CD20+ B cells and (E) CD20+ Bcl6+ Ki67+ GC B cells in the lymph node at 12 wpi are shown. Each data point represents one animal, and medians are indicated as horizontal lines. FDR adjusted p-values are reported in the graphs, FDR_p <0.05 was considered significant. See Table S4 for both unadjusted p and FDR_p for all comparisons.

### SHIV.C.CH505-infected infant and adult monkeys similarly develop virus tier 2 autologous plasma neutralization responses

Since the kinetics, magnitude, and breadth of plasma HIV Env-specific IgG responses were similar between infant and adult monkeys, we next evaluated the HIV neutralization activity of these plasma IgG responses. We found that both age groups developed similar neutralization activity at 12 wpi against the tier 1 clade-matched isolates MW965 (ID_50_ range infants: 135-31,736; adults: 324-4,949, p = 0.384) (Fig. 7A) and CH505 w4.3 (ID_50_ range infants: 45-950; adults: 45-750; p=0.605) (Fig. 7B). Further, neutralization activity against the autologous tier 2 neutralization sensitive challenge virus, CH505 T/F, was not observed in the majority of monkeys from both age groups until 12 wpi (Fig. 7C). Eight out of 12 adults (66.7%) and 3 out of 6 infants (50%) developed autologous virus neutralizing responses by 12wpi with no significant difference in potency (ID_50_ range infants: 45-161; adults: 45-380; p= 0.963) (Fig. 7C).

**Figure 7.**
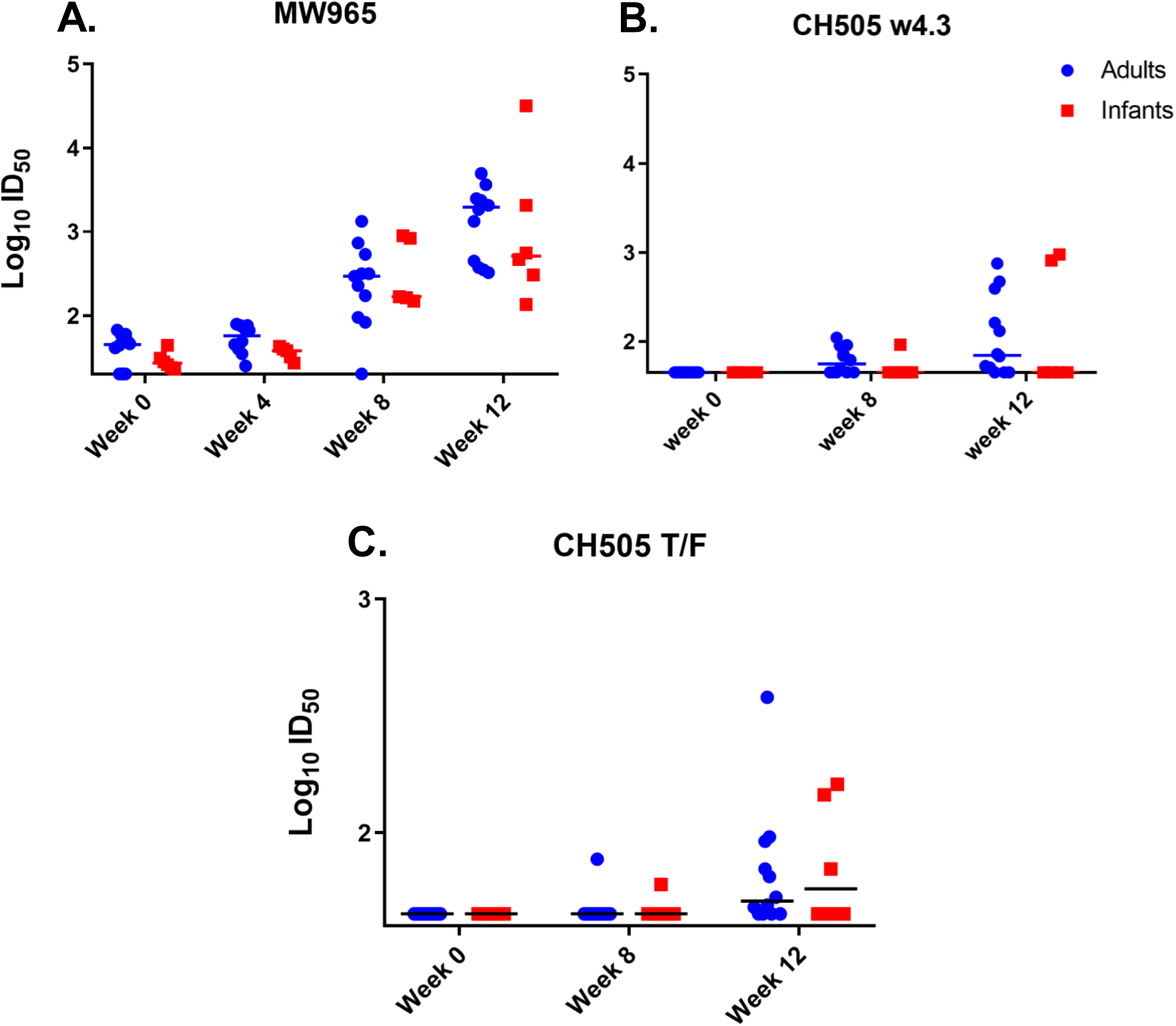
Magnitude and kinetics of plasma neutralization responses during acute SHIV.C.CH505 infection of infant and adult monkeys. The TZM-bl cell-based assay was performed to assess the neutralization activity of plasma antibodies. Tier 1 neutralization responses were evaluated against MW965 (A) and CH505 w4.3 (B) through 12 wpi. (C) Autologous virus neutralization titers against CH505 T/F. Each dot represents plasma neutralization of one monkey, and medians are indicated as horizontal lines.

### ADCC activity of plasma antibodies develops similarly in adult and infant acutely SHIV-infected monkeys

ADCC activity was assessed by the NK cell granzyme B response mediated by plasma from infant and adult RMs at 0, 6, and 10 wpi. ADCC activity was detectable in both age groups by week 6 of infection, and was maintained through week 10 with no significant difference in the magnitude or kinetics of the response between the two groups (6 wpi: p= 0.494 and 10 wpi: p= 0.591) (Fig. 8A). While similar kinetics were observed in CH505-specific ADCC Ab titers in adults and infants, infants exhibited plasma ADCC titers that were generally lower than adults, yet the differences were not statistically significant (6 wpi: p= 0.801 and 10 wpi: p= 0.807) (Fig. 8B).

**Figure 8.**
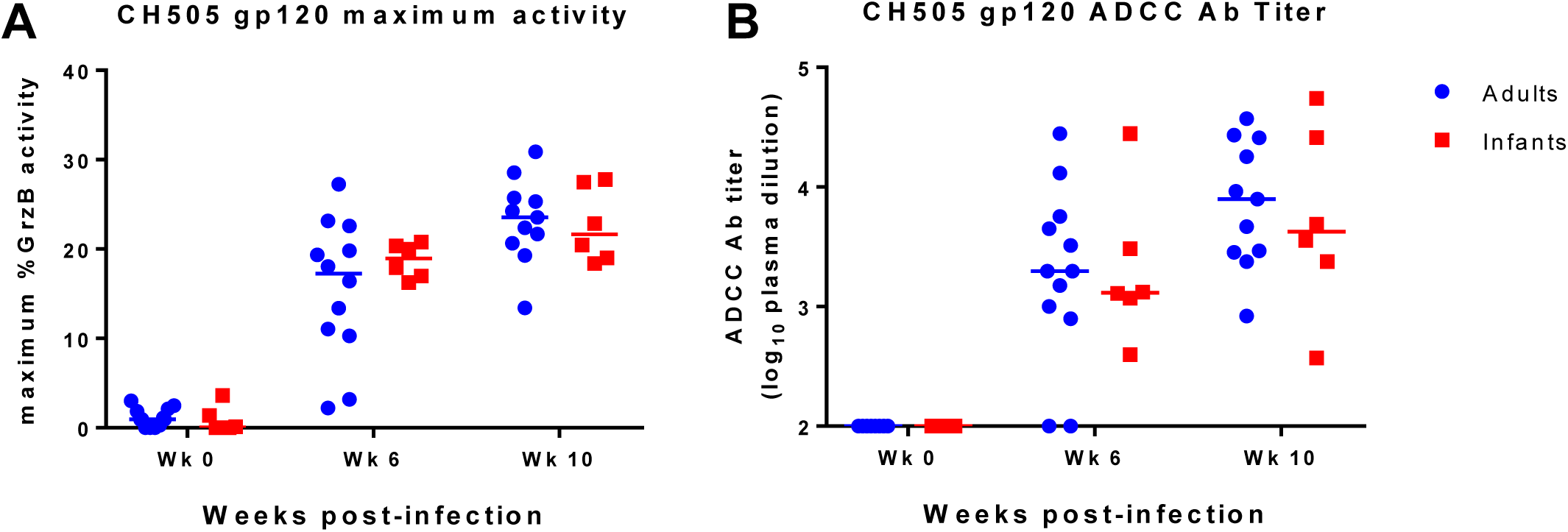
Similar ADCC activity of plasma antibodies of SHIV.C.CH505-infected infant and adult monkeys. ADCC activity was measured at weeks 0, 6, and 10 post-infection against CH505 gp120-coated target cells. The maximum granzyme B activity (A) and plasma dilution endpoint antibody titers (B) for each animal are shown. Medians are indicated as horizontal lines. See Table S3 for both unadjusted p and FDR_p for all comparisons

### Viral load at 12 wpi is significantly correlated with the development of autologous virus neutralization in infant and adult monkeys

We calculated spearman correlation coefficients to determine if a subset of the immunological responses assessed could predict the development of autologous neutralization in infant and adult monkeys at 12 wpi. CH505-specific gp120 IgG responses (rho=.69, p=.002, FDR p= .07) (Fig. 9B), ADCC antibody titers at 10 wpi (rho=.63, p=.01, FDR p=.16) (Fig. 9C), and CH505-specific Tfh cell frequencies at 12 wpi (rho=.49, p=.05, FDR p=.19) (Fig. 9D) were correlated with the development of autologous neutralization. However, these results were not statistically significant after adjustment for multiple comparisons. Yet, plasma viral load at 12 wpi was correlated with autologous neutralization after adjustment for multiple comparisons (rho=.76, p <.001, FDR p=.03) (Fig. 9A). A summary of all the immune parameters assessed can be found in Figure S3 and Table S5.

**Figure 9.**
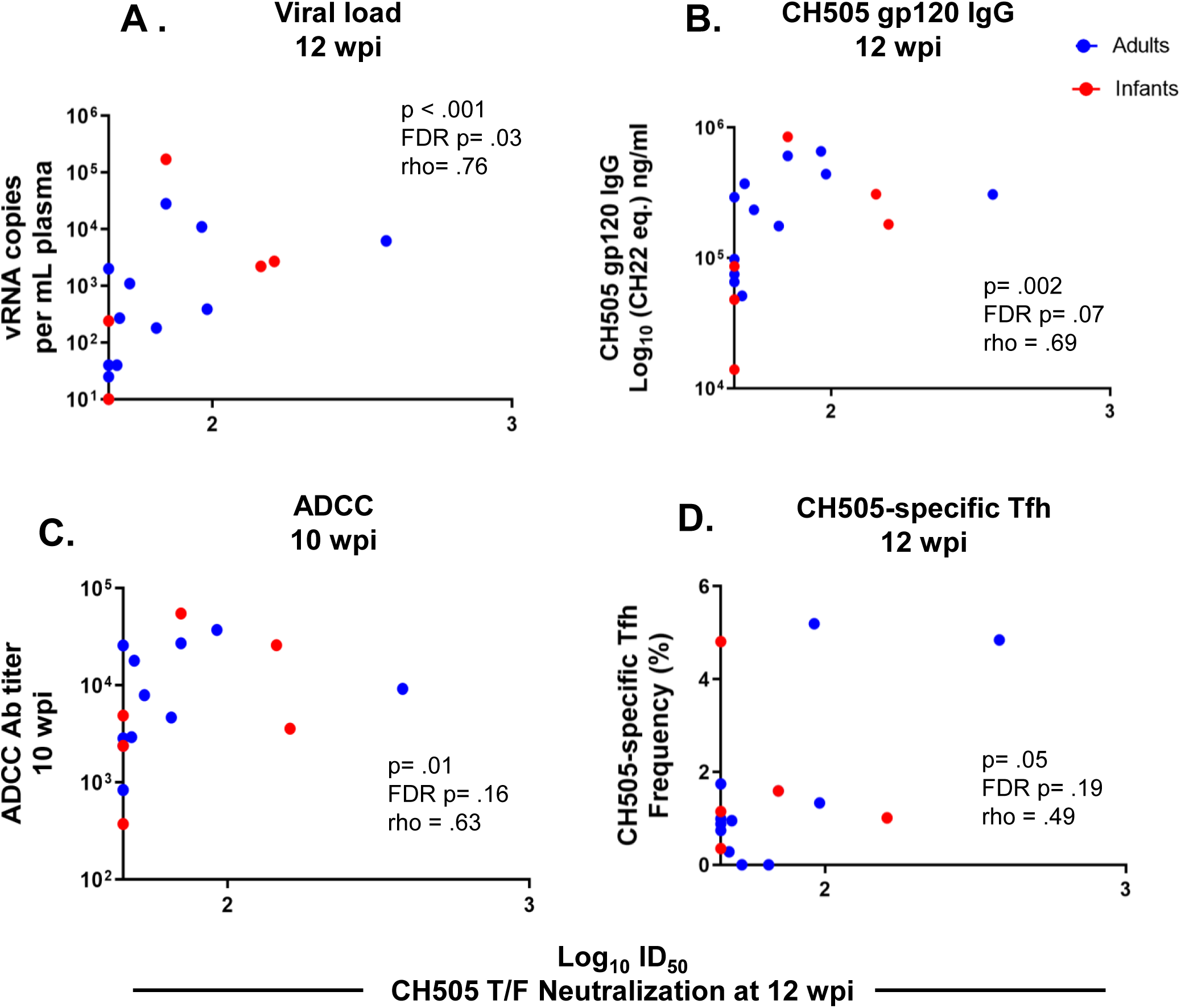
Viral load at 12 wpi is associated with the development of autologous neutralization. Correlations between CH505 T/F neutralization at 12 wpi and (A) plasma viral load at 12 wpi, (B) CH505 gp120 IgG at 12 wpi, (C) ADCC activity at 10 wpi, and (D) frequency of CH505-specific Tfh cells at 12 wpi. The coefficients of correlations (rho; ρ) and p values from testing whether the correlation coefiicent differed significantly from 0 are shown on the graphs. See Tables S5 and S6, and Figures S3 and S4 for a complete list of immune parameters tested and correlation coefficients with both unadjusted p and FDR_p.

ADCC activity has been suggested to be associated with reduced risk of infection and/or viral control in a number of studies (6,24, 25). We calculated spearman correlation coefficients to determine if a subset of the immunological parameters assessed were associated with the development antibodies capable of mediating ADCC. Overall, none of the immune responses evaluated appeared to predict ADCC activity. A summary of all the immune parameters assessed can be found in Figure S4 and Table S6.

## Discussion

The elimination of pediatric HIV infections and achievement of life-long immunity will likely require the development of successful immunization strategies tailored to the infant immune landscape. Thus, an understanding of the infant immune response and pathways for the development of neutralizing antibodies during HIV infection are critical to inform rational vaccine design. In this work, we utilized a rhesus macaque model of SHIV infection to better understand the development of HIV-specific immune responses in infants versus adults. The magnitude, kinetics, and specificity of HIV Env-specific plasma IgG responses were similar in SHIV.C.CH505 T/F-infected infant and adult monkeys. Furthermore, CH505 T/F-specific Tfh and memory B cell responses developed similarly in both age groups, consistent with the observed similarities in HIV Env-specific plasma IgG response. However, infant monkeys exhibited significantly higher frequencies of total Tfh and GC B cells in the lymph node during the early phase of infection. Moreover, acute SHIV.C.CH505 T/F infection elicited tier 2 autologous virus neutralization and ADCC responses that were similar in frequency and magnitude between both age groups. Lastly, correlation analysis determined that the magnitude of the plasma viral load was the strongest predictor of the development of autologous virus neutralization in both age groups.

A number of previous studies have demonstrated that differences exist between adult and pediatric immunity to HIV. A study of 46 HIV-infected human infants age 0 to 12 months suggests that infants develop antibodies against gp160 first, followed by anti-gp120 and –gp41 antibodies (11). However, the initial Env-specific antibody response in HIV-infected adults target gp41, and are non-neutralizing (10). While we observed similar kinetics of anti-gp120 plasma IgG antibodies in SHIV.C.CH505-infected infant and adult monkeys, gp41-specific responses exhibited a trend towards lower magnitude in the infant monkeys. Autologous virus neutralization responses also developed similarly in SHIV.C.CH505-infected adult and infant monkeys, with at least 50% of animals from both age groups exhibiting this response at 12 wpi (Fig. 7C). In human adults, autologous virus neutralization develops in approximately 3-6 months of infection (27-31), while neutralization breadth develops after 2-3 years (19, 20). In infants, exactly when autologous neutralizing antibodies develop is unknown and such analysis is difficult due to the presence of maternal antibodies. However, it has recently been recognized that HIV-infected infants can develop bnAbs as early as 1-2 years post-infection (13). If our results are reflective of what is happening in humans, it could suggest that the initial kinetics of neutralizing antibody responses is comparable between adults and infants, but subsequently infants acquire breadth faster than adults do. Interestingly, only plasma viral load was identified as a determinant of tier 2 autologous virus neutralization development among SHIV.C.CH505-infected infant and adult monkeys, suggesting that the development of autologous neutralization may be dependent upon antigen load. Further studies are needed to fully identify predictors of autologous neutralization, particularly since this response precedes the development of bnAbs in some individuals (20, 32, 33).

CD4+ T-follicular helper (Tfh) cells are crucial in providing help to B cells in the germinal center (GC) to support antibody maturation (34). A recent study of HIV-1 clade C-infected human children and adults demonstrated that frequencies of total Tfh (CXCR5+ PD1^hi^) and HIV-specific (Gag/Env), IL-21-producing GC-Tfh cells in oral lymphoid tissue were increased in older children receiving ART (age range: 6-10 years), however it is unknown whether this is true in early life (26). In our monkey model, we observed higher frequencies of total lymphoid Tfh cells and GC B cells in SHIV-infected infants compared to adults at 12 wpi (Fig. 5B and 6E). Yet, both age groups exhibited similar CH505-specifc Tfh frequencies at 12 wpi (Fig. 5A) corresponding with a similar magnitude in CH505 gp120 IgG binding responses (Fig. 2A) and systemic memory B cell responses (Fig. 6A and B). However, the quality of these Tfh responses may not be optimal early in infection, as we observed that only about 50% of the monkeys from both age groups developed autologous virus neutralization responses by 12 wpi. Tfh cells can further be characterized into distinct functional subsets, namely, Tfh1 (CXCR3+), Tfh2 (CXCR3-CCR6-), and Tfh17 (CCR6+) (35). Preferential enhancement of Tfh1 cells in the blood and lymph node have been observed in chronic SIV infection (36, 37), yet a recent study has reported the expansion of Tfh2 subsets during acute HIV infection in adults (38). We observed that during acute SHIV infection, adult monkeys exhibited significantly higher frequencies of Tfh1 cells where as Tfh17 cells were significantly higher in infants (Fig. 5C and E). Thus, acute SHIV infection induces a distinct Tfh phenotype in infants and adults. It is possible that distinctions in HIV immunity within our infant cohort are a result of maturation of the immune response rather than SHIV infection, thus future studies would need to include an age-matched infant control group. Nonetheless, imbalances in Tfh polarization may be implicated in HIV disease pathogenesis and further work in animal models are needed to define the mechanistic roles of Tfh phenotypes during HIV infection.

Our results suggest that the humoral immune response to SHIV infection develops similarly in adult and infant RMs, and corroborates with findings in human cohorts demonstrating that infants can develop robust HIV Env binding and neutralizing antibody responses despite their maturing immune landscape. Additionally, we have demonstrated that the Tfh landscape during acute infection in infants is distinct from that of adults, which may offer a potential advantage for infant vaccination, especially since studies suggest that infants are able to mount robust antibody responses to HIV Env vaccination, and these responses tend to be comparable or superior to that of adults (39-42). However, gaps in our knowledge still exists when it comes to understanding the infant immune response to HIV infection and how it can be harnessed for optimal vaccine-mediated protection against HIV-1. Thus, further development of infant SHIV models that depict human HIV-1 immunopathogenesis are imperative to further our understanding of infant HIV immunity and to inform vaccine elicitation of long-term protective immunity.

## Materials and Methods

### Animal care and sample collection

Adult female rhesus macaques ranged from 4 to 10 years of age, and infant rhesus macaques ranged from 6 weeks of age (Table 1). All macaques were of Indian origin, and from the type D retrovirus-free, SIV-free and STLV-1 free colony of the California National Primate Research Center (CNPRC; Davis, CA). Animals were maintained in accordance with the American Association for Accreditation of Laboratory Animal Care standards and The *Guide for the Care and Use of Laboratory Animals* (43). For sample collections, animals were sedated with ketamine HCl (Parke-Davis) injected at 10 mg/kg body weight. EDTA-anticoagulated blood was collected via peripheral venipuncture. Plasma was separated from whole blood by centrifugation, and PBMCs were isolated by density gradient centrifugation using Ficoll®-Paque (Sigma) or Lymphocyte Separation Medium (MP Biomedicals). All protocols were reviewed and approved by the University of California at Davis Institutional Animal Care and Use Committee (IACUC) prior to the initiation of the study.

### SHIV challenge of infant and adult monkeys

The generation of SHIV.C.CH505.375H.dCT has been previously described (20). The SHIV.C.CH505.375H.dCT challenge stock (provided by Dr. George M. Shaw, University of Pennsylvania) was prepared by infecting primary activated Indian rhesus macaque CD4+ T cells and 7-14 days later culture supernatants were pooled, as previously described (20). Virus titers were determined in TZM-bl cells, yielding 6.8×10^6^ TCID_50_ /ml.

Twelve adult monkeys were challenged intravenously with SHIV.C.CH505 at a dose of 3.4×10^5^ TCID_50_ (Table 1). Six infants were challenged orally beginning at 4 weeks of age. Initially, infants were exposed to SHIV.C.CH505 three times per day for 5 days with at a dose of 8.5×10^4^ TCID_50_/ml in an isotonic sucrose solution and bottle-fed, in order to simulate oral acquisition via breastfeeding. After one week, only one infant became infected, and the remaining five infants were challenged three weeks later under sedation at a dose of 6.8×10^5^ TCID_50_/ml until infected. After three weeks one infant remain uninfected and thus was subsequently challenged at an increasing dose (3.4×10^6^ TCID_50_/ml) until infected (Table 1).

### Viral RNA load quantification

Plasma RNA load was quantified using a well-established quantitative reverse transcriptase (RT) PCR assay targeting SIVgag RNA, as previously described (18). RNA was isolated from plasma samples using the QIAsymphony Virus/Bacteria Midi kit on the QIAsymphony SP automated sample preparation platform (Qiagen, Hilden, Germany). RNA was extracted manually if plasma volumes were limited. Data reported are the number of SIV RNA copy equivalents per ml of plasma, with a limit of detection of 15 copies/ ml.

### Lymphoctye counts

Absolute lymphocyte counts in blood were calculated using the PBMC counts obtained by automated complete blood counts, multiplied by the lymphocyte percentages.

### Enzyme-Linked Immunosorbent Assay (ELISA), recombinant protein and soluble CD4 blocking

Env-binding IgG was assessed in plasma in a 384-well plate format. The plates were coated overnight with HIV CH505 gp120 (30 ng/well) or MN gp41 (3 µg/ml) and then blocked with the assay diluent (phosphate-buffered saline containing 4% whey, 15% normal goat serum, and 0.5% Tween 20). Serially diluted plasma were then added to the plates and incubated for 1 hour, followed by detection with a horseradish peroxidase (HRP)-conjugated antibody, polyclonal goat anti-monkey IgG (Rockland Immunochemicals). The plates were developed by using the ABTS-2 peroxidase substrate system (KPL). The monoclonal antibody, b12R1, was used to develop standard curves, and the concentration of IgG antibody was calculated relative to the standard using a 5-parameter fit curve (SoftMax Pro 7). For monoclonal antibodies, effective concentration 50% (EC50) was calculated by the concentration of antibody which resulted in a 50% reduction in optical density (OD) from the maximum value.

For CD4 blocking ELISAs, 384-well plates (Corning Life Sciences) were coated with C.1086 gp120 at 30ng/well. Following the same steps as previously stated, plates were blocked with assay diluent and serially diluted monoclonal antibody and plasma were added, and incubated for 1 hour. Soluble CD4 (sCD4) (NIH AIDS Reagent Program, Division of AIDS, NIAID, NIH: Human Soluble CD4 Recombinant Protein (sCD4) from Progenics) was then added at concentration of 0.64 µg/mL. The sCD4 binding was detected using a biotinylated Human anti-CD4 (Thermo Fisher Scientific) followed by HRP-conjugated Streptavidin. Percent sCD4 binding inhibition was calculated as follows: 100 – (average of sera duplicate OD/average of negative control OD) x 100. OD referring to optical density. A reduction of absorbance by >50% by Abs present in plasma indicated blocking of sCD4 binding to C.1086 gp120.

### Binding Antibody Multiplex Assay (BAMA)

HIV-1 epitope specificity and breadth were determined using BAMA, as previously described (10). HIV-1 antigens were conjugated to polystyrene beads (Bio-Rad) as previously described (18), then binding of IgG to the bead-conjugated HIV-1 antigens was measured in plasma samples from the infant and adult monkey cohorts. The positive control was purified IgG from a pooled plasma of HIV Env-vaccinated rhesus macaques (RIVIG) (44). The conjugated beads were incubated on filter plates (Millipore) for approximately 30 minutes before plasma samples were added. The plasma samples were diluted in assay diluent (1% dry milk + 5% goat serum + 0.05% tween-20 in 1X phosphate buffer saline, pH 7.4.) at a 1:500-point dilution. Beads and diluted samples were incubated for 30 minutes, then IgG binding was detected using a PE-conjugated mouse anti-monkey IgG (Southern Biotech) at 4 μg/mL. The beads were washed and acquired on a Bio-Plex 200 instrument (Bio-Rad) and IgG binding was expressed as mean fluorescence intensity (MFI). To assess assay background, the MFI of binding to wells that did not contain beads or sample (blank wells) and non-specific binding of the samples to unconjugated blank beads were evaluated during assay analysis. High background detection for plasma samples were noted and repeated if necessary. An HIV-envelope specific antibody response was considered positive if above the lower limit of detection (100 MFI). To check for consistency between assays, the EC50 and maximum MFI values of the positive control (RIVIG) was tracked by Levy-Jennings charts. The antigens conjugated to the polystyrene beads are as follows: C. 1086 gp140, C.1086 gp120, A1.Con_env03 gp140, A233 gp120, B.Con_env03 gp140, Con6 gp120, ConC gp120, MN gp120, Linear V2. B, V3.C, C5.2.C, C1, conformational V1V2, ConC V3, and MN V3, and C. 1086 V1V2 (Table S1).

### Linear peptide microarray mapping and data analysis

Solid phase peptide microarray epitope mapping was performed as previously described (45), with minor modifications. Briefly, array JPT Peptide Technologies GmbH (Germany) prepare arrays slides by printing a library designed by Dr. B. Korber, Los Alamos National Laboratory, onto Epoxy glass slides (PolyAn GmbH, Germany). The library contains 15-mer peptides overlapping by 12, covering consensus Env (gp160) clade A, B, C, D, Group M, CRF1, and CRF2 and vaccine strains (gp120) 1.A244, 1.TH023, MN, C.1086, C.TV1, and C.ZM651. To assess CH505-specific responses, a peptide library containing 59 CH505 strains (gp120, gp145, gp160, and SOSIP, sequences provided by Dr. Barton Haynes, Duke University) was also designed (20). Sera were diluted 1/50 and applied to the peptide array, followed by washing and detection using goat anti-human IgG-Alexa Fluor 647. Array slides were scanned at a wavelength of 635 nm with an InnoScan 710 AL scanner (Innopsys, France) using XDR mode. Scan images were analyzed using MagPix 8.0 software to obtain binding intensity values for all peptides. Microarray data were then processing using R package pepStat (46) to obtain binding signal for each peptide, which is defined as log_2_(Intensity of 12 wpi sample/intensity of matched baseline sample). Binding magnitude to each identified epitope is defined as the highest binding signal by a single peptide within the epitope region.

### Neutralization assays

Neutralization by antibodies in plasma of MW965.LucR.T2A.ecto/293T IMC (clade C, tier 1), CH505 w4.3 HIV-1 pseudovirus (clade C, tier 1a), and autologous CH505.TF (clade C, tier 2) HIV-1 pseudovirus was measured in TZM-bl cells (NIH AIDS Reagent Program, Division of AIDS, NIAID, NIH; from John Kappes) via a reduction in luciferase reporter gene expression after a single round of infection as previously described (47-49). Prior to screening, plasma was heated-inactivated at 56°C for 30 min. Luminescence was measured using a Victor X3 multilabel plate reader, 1 s per well (PerkinElmer). The ID_50_ was calculated as the dilution that resulted in a 50% reduction in relative luminescence units (RLU) compared to virus control wells. The monoclonal antibody, b12R1, was used as a positive control for MW965 assays, and VRC01 was used a positive control for all other assays.

### ADCC

The ADCC-GTL assay was used to measure plasma ADCC activity as previously described (50). Briefly, CEM.NKRCCR5 target cells (NIH AIDS Reagent Program, Division of AIDS, NIAID, NIH; from Alexandra Trkola) (51) were coated with recombinant CH505 or 1086.C K160N gp120. Cryopreserved human peripheral blood mononuclear cells (PBMCs) from an HIV-1 seronegative donor with the heterozygous 158 F/V genotype for the Fcγ receptor IIIa were used as the source of effector cells (52, 53). Adult and infant plasma samples were tested after a 4-fold serial dilution starting at 1:100. ADCC was measured as percent Granzyme B (GzB) activity, defined as the frequency of target cells positive for proteolytically active GzB out of the total viable target cell population. Final results are expressed after subtraction of the background GzB activity observed in wells containing target and effector cells in the absence of plasma. ADCC endpoint titers were determined by interpolating the last positive dilution of plasma (>8% GzB activity).

### CH505 envelope-specific memory B cell phenotyping

For phenotyping of CH505 Env-specific memory B cells, suspension of 10^6^ PBMCs were blocked with 6.25 μg/ml anti-human CD4 antibody (BD Biosciences) at 4°C for 15 min. After incubation, PBMCs were washed twice with PBS, and pelleted at 1500 rpm for 5 min. PBMCs were then incubated at 4°C with LIVE/DEAD Fixable Aqua Dead Cell Stain Kit (Thermo Fisher Scientific) for 30 minutes. Following incubation and wash with PBS, PBMCs were then stained with a cocktail of fluorescently conjugated antibodies for surface markers including CD20, CD3, IgM, CD16, CD8, IgD PE, CD14 and CD27 (Table S2) and custom-conjugated BV421-HIV-1 gp120 (C.CH505 T/F) and AF647-HIV-1 gp120 (C.CH505 T/F) prepared as described previously (54). The stained PBMCs were acquired on an LSRII flow cytometer (BD Biosciences) using BD FACS Diva software, and analyzed with FlowJo software version 10. The following gating strategy was applied: lymphocytes were gated on singlets and live cells were selected to gate on CD3^−^cells (T cells), CD14^−^cells (monocytes/macrophages), and CD20^+^cells (B cells). B cells were further gated on CD27^+^memory B cells. Only B cells positive for both BV421-gp120 and AF647-gp120 were considered CH505-Env specific. For a detailed list of antibodies used for B cell phenotyping, see Table S2.

### Lymph node Tfh phenotyping and Activation-Induced Marker (AIM) Assay

The AIM assay was based on previous work (55). Briefly, cryopreserved rhesus macaque lymph node cells (12 wpi) were thawed, rested for 3h at 37°C/5% CO_2_, re-suspended in AIM V medium (Gibco), and transferred to wells of a 24-well plate at 10^6^ cells per well. Cells were cultured for 18 hr at 37°C/5% CO_2_ with no exogenous stimulation or with gp140 stimulation (5 µg/ml CH505 T/F gp140 protein and 0.5 µg/ml of a 15-mer peptide pool with 11-residue overlap spanning CH505 T/F gp140). As a positive control, cells were stimulated with 0.5 µg/ml Staphylococcal enterotoxin B (SEB) (Sigma). Duplicates of each condition were performed when cell numbers permitted. Following stimulation, cells were labeled with fluorescently labelled antibodies to the following surface antigens: PD-1, CD8a, CD25, CD4, CD20, CD69, CD137, CD196, OX40, CD183, CD3, CD45RA, and CD185. For Tfh phenotyping the following gating strategy was applied: lymphocytes were gated on singlets and live cells were selected to gate on CD4+ CXCR5+ Foxp3-cells, followed by gating on CCR6 and CXCR3. Cell viability was measured using Live/Dead Fix Aqua stain (eBioscience). Flow cytometry data were acquired on a LSRII running FACSDiva software (BD Biosciences) and analyzed on FlowJo (FlowJo). For a detailed list of antibodies used for phenotyping see Table S2, and for the AIM assay gating strategy see Figure S2.

## Statistical Methods

Immune assay measurements at various time points post-infection and the change in immune assay measurements from baseline were compared between SHIV-infected infant and adult monkeys using Wilcoxon rank sum tests with exact p-values. Spearman’s rank correlation coefficients were estimated for the cohort as a whole as well as by adult monkeys and infants separately. All correlations were tested with exact p-values to assess whether any were significantly different from zero. To adjust for multiple comparisons, the Benjamini–Hochberg (BH) procedure was used to control the false discovery rate (FDR). Separate adjustments to control the FDR at α = 0.05 were performed for comparisons between the infant and adult monkeys using: 1) the pre-specified primary endpoints for a total of 26 tests (Table S3); 2) the pre-specified secondary endpoints for a total of 43 tests (Table S4); 3) correlations between pre-specified parameters and CH505.TF neutralization at 12 wpi for a total of 60 tests (Table S5); and 4) correlations between pre-specified parameters and ADCC - %Grz B activity at 10 weeks post-infection for a total of 15 tests (Table S6). Both the unadjusted (raw_p) and FDR-adjusted (FDR_p) p-values are reported in Tables S3-S6. All statistical tests were performed using SAS version 9.4 (Cary, NC, USA).

## Acknowledgements

The work was supported by National Institutes of Health grants P01 AI117915 (S.R.P and K.D.P); T32 CA911139 (A.N.N); 5R01 AI106380 (S.R.P); and 5R01-DE025444 (S.R.P). This work was also supported by the Penn Center for AIDS Research Viral and Molecular Core P30 AI045008 (K.B.); the BEAT-HIV: Delaney Collaboratory to Cure HIV-1 Infection by Combination Immunotherapy UM1AI126620 (K.B.); and CARE: Delaney Collaboratory for AIDS Eradication UM1AI126619 (K.B.).

We would like to thank Dr. George Shaw, University of Pennsylvania Department of Medicine, Philadelphia, PA for providing us with SHIV.C.CH505.375H.dCT. Protein antigens for BAMAs and ELISAs were generously provided by Kevin Saunders and Barton Haynes, supported by NIH NIAID Division of AIDS UM1 grantAI100645 for the Center for HIV/AIDS Vaccine Immunology-Immunogen Discovery (CHAVI-ID), and produced at the DHVI Protein Production Facility. Study data were collected and managed using REDCap (Research Electronic Data Capture) electronic data capture tools hosted at Duke University.

The authors thank Jeff Lifson, Rebecca Shoemaker and colleagues in the Quantitative Molecular Diagnostics Core of the AIDS and Cancer Virus Program of the Frederick National Laboratory for expert assistance with viral load measurements. We would also like to thank J. Watanabe, J. Usachenko, A. Ardeshir and the staff of the CNPRC Colony Research Services for their support in these studies. We thank Dr. Georgia Tomaras’ laboratory for technical support with linear epitope mapping microarray, and R. Whitney Edwards for performing ADCC assays. Flow cytometry was performed in the Duke Human Vaccine Institute Flow Cytometry Facility (Durham, NC). We are also thankful for statistical support for this study provided by the Center for AIDS Research at the University of North Carolina at Chapel Hill, an NIH-funded program (P30 AI50410). The funders had no role in study design, data collection and interpretation, or the decision to submit the work for publication. The content is solely the responsibility of the authors and does not necessarily represent the official views of the National Institutes of Health.

